# Gene model for the ortholog of *Ilp3* in *Drosophila eugracilis*

**DOI:** 10.1101/2025.08.16.670678

**Authors:** James O’Brien, Ryan Dufur, Martina Kotrba, Jeremiah Severn, Jacqueline Wittke-Thompson, Chinmay Rele, Christine M. Fleet

**Affiliations:** Oklahoma Christian University, Edmond, OK; University of St. Francis, Joliet, Illinois; Emory & Henry University, Emory, VA; University of Alabama, Tuscaloosa, AL, USA

## Abstract

Gene model for the ortholog of *Insulin-like peptide 3* (*Ilp3*) in the D. eugracilis Apr. 2013 (BCM-HGSC/Deug_2.0) (DeugGB2) Genome Assembly of *D. eugracilis* (GenBank Accession: GCA_000236325.2) of *Drosophila eugracilis*. This ortholog was characterized as part of a dataset to study the evolution of the Insulin/ insulin-like growth factor signaling pathway (IIS) across the genus Drosophila using the Genomics Education Partnership gene annotation protocol for Course-based Undergraduate Research Experiences.

## Introduction

*This article reports a predicted gene model generated by undergraduate work using a structured gene model annotation protocol defined by the Genomics Education Partnership (GEP; thegep.org) for Course-based Undergraduate Research Experience (CURE). The following information in quotes may be repeated in other articles submitted by participants using the same GEP CURE protocol for annotating Drosophila species orthologs of* Drosophila melanogaster *genes in the insulin signaling pathway*.

“Computational gene predictions in non-model organisms often can be improved by careful manual annotation and curation, allowing for more accurate analyses of gene and genome evolution (Mudge and Harrow 2016; Tello-Ruiz et al., 2019). The Genomics Education Partnership (thegep.org) uses web-based tools to allow undergraduates to participate in course-based research by generating manual annotations of genes in non-model species (Rele et al., 2023). These models of orthologous genes across species, such as the one presented here, then provide a reliable basis for further evolutionary genomic analyses when made available to the scientific community. The particular gene ortholog described here, Insulin-like peptide 3 (Ilp3) in D. eugracilis, was characterized as part of a developing dataset to study the evolution of the Insulin/insulin-like growth factor signaling pathway (IIS) across the genus Drosophila.” (Myers et al., 2024).

“The IIS pathway is a highly conserved signaling pathway in animals and is central to mediating organismal responses to nutrients (Hietakangas and Cohen 2009; Grewal 2009)” (Myers et al., 2024). “Invertebrate insulins function similarly to metazoan insulin-like growth factors and play a role in cell and organ growth (Chan 2000). In *Drosophila*, seven insulin-like peptides (Ilp1-Ilp7) have a two-chain structure similar to vertebrate insulin and interact with the sole insulin-like receptor, InR, to initiate the insulin signaling cascade (Brogiolo et al., 2001; Nässel and Broeck 2016). Like the *Ilp2* and *Ilp5* genes, the *Ilp3* gene is expressed in median neurosecretory cells (MNCs) in the brain (Ikeya et al., 2002). While the seven Ilps act redundantly with respect to promoting growth, they also have unique expression patterns and functions (Ikeya et al., 2002; Grönke et al., 2010). Ilp3 may act with the transcription factor dFOXO in a positive feedback loop to regulate Ilp2 and Ilp5 secretion from MNCs (Grönke et al., 2010). In female *Drosophila*, ablation of MNCs or knockout of *Ilp3* have been shown to reduce fecundity and remating rates (Grönke et al., 2010; Wigby et al., 2011). Knockout of *Ilp3* also results in sleep defects (Yamaguchi et al., 2022).” (Gruys et al., 2024).

“*D. eugracilis* (NCBI taxon ID 29029) is part of the *melanogaste*r species group within the subgenus *Sophophora* of the genus *Drosophila* (Pélandakis and Solignac, 1993). It was first described as *Tanygastrella gracilis* by Duda (1924) and revised to *Drosophila eugracilis* by Bock and Wheeler (1972). *D. eugracilis* is found in humid tropical and subtropical forests across southeast Asia (https://www.taxodros.uzh.ch, accessed 1 Feb 2023).” (Morgan et al., 2022).

We propose a gene model for the *D. eugracilis* ortholog of the *D. melanogaster Insulin-like peptide 3* (*Ilp3*) gene. The genomic region of the ortholog corresponds to the uncharacterized protein XP_017080989.1 (Locus ID LOC108114482) in the D. eugracilis Apr. 2013 (BCM-HGSC/Deug_2.0) (DeugGB2) Genome Assembly of *D*. eugracilis (GCA_000236325.2 - Chen et al., 2014). This model is based on RNA-Seq data from *D. eugracilis* (PRJNA63469) and *Ilp3* in *D. melanogaster* using FlyBase release FB2024_02 (GCA_000001215.4; Gramates et al., 2022; Jenkins et al., 2022; Larkin et al., 2021).

### Synteny

The target gene, *Ilp3*, occurs on chromosome 3L in *D. melanogaster* and is nested by *CG32052* alongside *Insulin-like peptide 4* (*Ilp4*) (upstream) and *Insulin-like peptide 2* (*Ilp2*) (downstream). *Ilp3* is flanked further upstream by *Inhibitor-2* (*I-2*) and CG43897, which nests *Insulin-like peptide 5* (*Ilp5*) and downstream by *Insulin-like peptide 1* (*Ilp1*) and *Z band alternatively spliced PDZ-motif protein 67* (*Zasp67*). The *tblastn* search of *D. melanogaster* Ilp3-PA (query) against the *D. eugracilis* Apr. 2013 (BCM-HGSC/Deug_2.0) (DeugGB2) Genome Assembly of *D. eugracilis* (GCA_000236325.2 - Chen et al., 2014) placed the putative ortholog of Ilp3 within scaffold scf7180000409711 (KB465257.1) at locus LOC108114482 (XP_017080989.1)— with an E-value of 7e-16 and a percent identity of 46.43%. Furthermore, the putative ortholog is nested by LOC108114479 (XP_017080986.1) alongside LOC108114483 (XP_017080990.1) upstream, and LOC108113894 (XP_017080088.1) and LOC108114481 (XP_017080988.1) downstream (E-value: 0.0, 1e-38, 1e-68 and 3e-53; identity: 91.26%, 60.00%, 57.40% and 59.71%, respectively, as determined by *blastp*; Figure 1A, Altschul et al., 1990). The putative ortholog is flanked further upstream by LOC108114226 (XP_017080582.1) and LOC108114224 (XP_017080566.1), which nests LOC108114228 (XP_017080583.1); that correspond to *I-2*, CG43897 and *Ilp5* in *D. melanogaster* (E-value: 1e-100, 0.0 and 2e-38; identity: 85.37%, 80.83% and 56.25%, respectively, as determined by *blastp*). The putative ortholog of *Ilp3* is flanked downstream by LOC108114480 (XP_017080987.1) and LOC108114478 (XP_017080982.1), which correspond to *Ilp1* and *Zasp67* in *D. melanogaster* (E-value: 3e-59 and 0.0; identity: 63.64% and 82.66%, respectively, as determined by *blastp*). The putative ortholog assignment for *Ilp3* in *D. eugracilis* is supported by the following evidence: The genes surrounding the *Ilp3* ortholog are orthologous to the genes at the same locus in *D. melanogaster*, aside from the insertion of *CG33483*. Local synteny is completely conserved, supported by results generated from *blastp*, so we conclude that LOC108114482 is the correct ortholog of *Ilp3* in *D. eugracilis* (Figure 1A).

**Figure 1.**
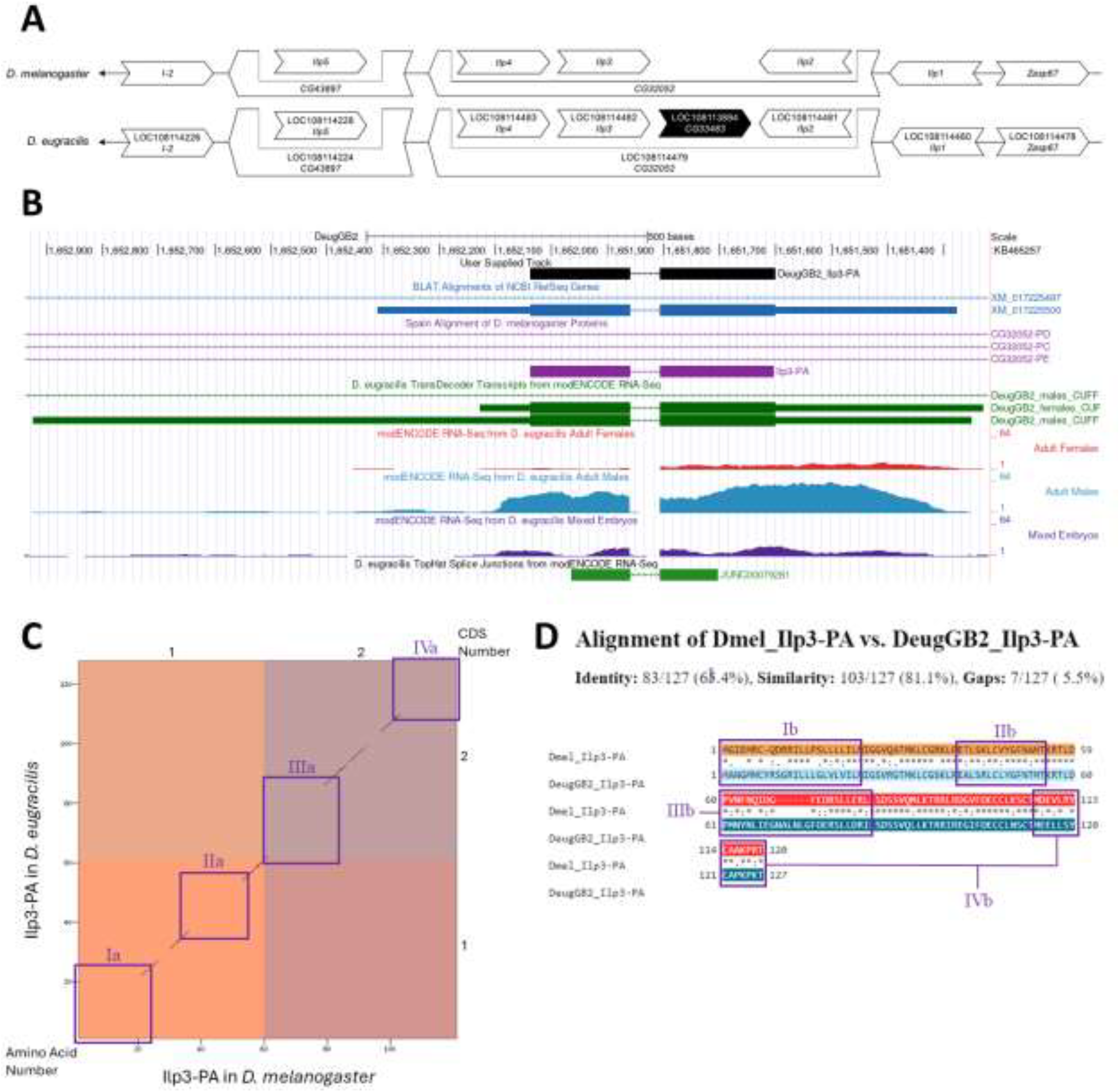
*Ilp3* gene model comparison between *Drosophila eugracilis* and *Drosophila melanogaster* orthologs. **(A) Synteny comparison of the genomic neighborhoods for *Ilp3* in *Drosophila melanogaster* and *D. eugracilis***. Thin underlying arrows indicate the DNA strand within which the target gene–*Ilp3*–is located in *D. melanogaster* (top) and *D. eugracilis* (bottom). Thin arrows pointing to the left indicate that *Ilp3* is on the negative (-) strand in *D. eugracilis* and *D. melanogaster*. The wide gene arrows pointing in the same direction as *Ilp3* are on the same strand relative to the thin underlying arrows, while wide gene arrows pointing in the opposite direction of *Ilp3* are on the opposite strand relative to the thin underlying arrows. White gene arrows in *D. eugracilis* indicate orthology to the corresponding gene in *D. melanogaster*, while black gene arrows indicate non-orthology. Gene symbols given in the *D. eugracilis* gene arrows indicate the orthologous gene in *D. melanogaster*, while the locus identifiers are specific to *D. eugracilis*. **(B) Gene Model in GEP UCSC Track Data Hub (Raney et al**., **2014)**. The coding-regions of *Ilp3* in *D. eugracilis* are displayed in the User Supplied Track (black); coding CDSs are depicted by thick rectangles and introns by thin lines with arrows indicating the direction of transcription. Subsequent evidence tracks include BLAT Alignments of NCBI RefSeq Genes (dark blue, alignment of Ref-Seq genes for *D. eugracilis*), Spaln of D. melanogaster Proteins (purple, alignment of Ref-Seq proteins from *D. melanogaster*), Transcripts and Coding Regions Predicted by TransDecoder (dark green), RNA-Seq from Adult Females, Adult Males and Mixed Embryos (red, light blue and purple, respectively; alignment of Illumina RNA-Seq reads from *D. eugracilis*), and Splice Junctions Predicted by regtools using *D. eugracilis* RNA-Seq (PRJNA63469). The splice junction shown in green (JUNC00079281) has a read-depth of 50. **(C) Dot Plot of Ilp3-PA in *D. melanogaster* (*x*-axis) vs. the orthologous peptide in *D. eugracilis* (*y*-axis)**. Amino acid number is indicated along the left and bottom; CDS number is indicated along the top and right, and CDSs are also highlighted with alternating colors. Line breaks in the dot plot indicate mismatching amino acids at the specified location between species. Regions that lack conservation are highlighted in purple (Box Ia, IIa, IIIa and IVa, respectively). **(D) Protein alignment between *D. melanogaster* Ilp3-PA and its putative ortholog in *D. eugracilis***. The alternating colored rectangles represent adjacent exons. The symbols in the match line denote the level of similarity between the aligned residues. An asterisk (*) indicates that the aligned residues are identical. A colon (:) indicates the aligned residues have highly similar chemical properties—roughly equivalent to scoring > 0.5 in the Gonnet PAM 250 matrix (Gonnet et al., 1992). A period (.) indicates that the aligned residues have weakly similar chemically properties—roughly equivalent to scoring > 0 and ≤ 0.5 in the Gonnet PAM 250 matrix. A space indicates a gap or mismatch when the aligned residues have a complete lack of similarity—roughly equivalent to scoring ≤ 0 in the Gonnet PAM 250 matrix. Areas highlighted in purple (Ib, IIb, IIIb and IVb) correspond to similarly labeled regions in the dot plot (Ia, IIa, IIIa and IVa).

### Protein Model

*Ilp3* in *D. eugracilis* has two CDSs within the genome sequence. The unique protein sequence (Ilp3-PA) is translated from one mRNA isoform (*Ilp3-RA*; Figure 1B). Relative to the ortholog in *D. melanogaster*, the CDS number and protein isoform count are conserved. The sequence of Ilp3-PA in *D. eugracilis* has 68.63% identity (E-value: 2e-48) with the protein-coding isoform Ilp3-PA in *D. melanogaster*, as determined by *blastp* (Figure 1C). Regions which lack conservation are highlighted in purple in the dot plot and protein alignment (I, II, III and IV in Figure 1C and Figure 1D, respectively). Coordinates of this curated gene model are stored by NCBI at GenBank/BankIt (accession **<u>BK059552</u>)**. This gene model can also be seen within the target genome at this [TrackHub](https://gander.wustl.edu/cgi-bin/hgTracks?db=DeugGB2&lastVirtModeType=default&lastVirtModeExtraState=&virtModeType=default&virtMode=0&nonVirtPosition=&position=KB465257%3A1651102%2D1652538&hgsid=16740525_wQgraWPI6jl46UsiaoKRLWPlag4v).

## Methods

“Detailed methods including algorithms, database versions, and citations for the complete annotation process can be found in Rele et al. (2023). Briefly, students use the GEP instance of the UCSC Genome Browser v.435 (https://gander.wustl.edu; Kent WJ et al., 2002; Navarro Gonzalez et al., 2021) to examine the genomic neighborhood of their reference IIS gene in the D. melanogaster genome assembly (Aug. 2014; BDGP Release 6 + ISO1 MT/dm6). Students then retrieve the protein sequence for the D. melanogaster reference gene for a given isoform and run it using tblastn against their target Drosophila species genome assembly on the NCBI BLAST server (https://blast.ncbi.nlm.nih.gov/Blast.cgi; Altschul et al., 1990) to identify potential orthologs. To validate the potential ortholog, students compare the local genomic neighborhood of their potential ortholog with the genomic neighborhood of their reference gene in D. melanogaster. This local synteny analysis includes at minimum the two upstream and downstream genes relative to their putative ortholog. They also explore other sets of genomic evidence using multiple alignment tracks in the Genome Browser, including BLAT alignments of RefSeq Genes, Spaln alignment of D. melanogaster proteins, multiple gene prediction tracks (e.g., GeMoMa, Geneid, Augustus), and modENCODE RNA-Seq from the target species. Detailed explanation of how these lines of genomic evidenced are leveraged by students in gene model development are described in Rele et al. (2023). Genomic structure information (e.g., CDSs, intron-exon number and boundaries, number of isoforms) for the D. melanogaster reference gene is retrieved through the Gene Record Finder (https://gander.wustl.edu/~wilson/dmelgenerecord/index.html; Rele et al., 2023). Approximate splice sites within the target gene are determined using tblastn using the CDSs from the D. melanogaster reference gene. Coordinates of CDSs are then refined by examining aligned modENCODE RNA-Seq data, and by applying paradigms of molecular biology such as identifying canonical splice site sequences and ensuring the maintenance of an open reading frame across hypothesized splice sites. Students then confirm the biological validity of their target gene model using the Gene Model Checker (https://gander.wustl.edu/~wilson/dmelgenerecord/index.html; Rele et al., 2023), which compares the structure and translated sequence from their hypothesized target gene model against the D. melanogaster reference gene model. At least two independent models for a gene are generated by students under mentorship of their faculty course instructors. Those models are then reconciled by a third independent researcher mentored by the project leaders to produce the final model. Note: comparison of 5’ and 3’ UTR sequence information is not included in this GEP CURE protocol.” (Gruys et al., 2025)

## Supplemental files

1. Zip file containing a FASTA, PEP, GFF files for the gene model
2. Figure 1 in high resolution

## Metadata

Bioinformatics, Genomics, *Drosophila*, Genotype Data, New Finding

## Acknowledgements

We would like to thank Wilson Leung for developing and maintaining the technological infrastructure that was used to create this gene model, Madeline L. Gruys for retrofitting this model and Laura K. Reed for overseeing the project. Thank you to FlyBase for providing the definitive database for *Drosophila melanogaster* gene models. Further, we would like to thank the editors and developers at the journal *microPublication: Biology* for assistance in developing the template for these single gene ortholog publications.

## Funding

This material is based upon work supported by the National Science Foundation (1915544) and the National Institute of General Medical Sciences of the National Institutes of Health (R25GM130517) to the Genomics Education Partnership (GEP; https://thegep.org/; PI-LKR). Any opinions, findings, and conclusions or recommendations expressed in this material are solely those of the author(s) and do not necessarily reflect the official views of the National Science Foundation nor the National Institutes of Health.

## Notes

https://gander.wustl.edu/cgi-bin/hgTracks?db=DeugGB2&lastVirtModeType=default&lastVirtModeExtraState=&virtModeType=default&virtMode=0&nonVirtPosition=&position=KB465257%3A1651102%2D1652538&hgsid=16740525_wQgraWPI6jl46UsiaoKRLWPlag4v

### Competing Interest Statement

The authors have declared no competing interest.

## References

Altschul, S.F., Gish, W., Miller, W., Myers, E.W. & Lipman, D.J. 1990. “Basic local alignment search tool.” J. Mol. Biol. 215:403–410. PMID: 2231712

Bock IR, Wheeler MR. 1972. The Drosophila melanogaster species group. Univ. Texas Publs Stud. Genet. 7(7213):1-102. FBrf0024428

Brogiolo W, Stocker H, Ikeya T, Rintelen F, Fernandez R, Hafen E. 2001. An evolutionarily conserved function of the Drosophila insulin receptor and insulin-like peptides in growth control. Curr Biol. 11(4):213–221. PMID: 11250149

Chan SJ, Steiner DF. 2000. Insulin Through the Ages: Phylogeny of a Growth Promoting and Metabolic Regulatory Hormone. Integ Comp Biol. 40(2):213–222. doi.org/10.1093/icb/40.2.213

Chen ZX, Sturgill D, Qu J, Jiang H, Park S, Boley N, Suzuki AM, Fletcher AR, Plachetzki DC, FitzGerald PC, Artieri CG, Atallah J, Barmina O, Brown JB, Blankenburg KP, Clough E, Dasgupta A, Gubbala S, Han Y, Jayaseelan JC, Kalra D, Kim YA, Kovar CL, Lee SL, Li M, Malley JD, Malone JH, Mathew T, Mattiuzzo NR, Munidasa M, Muzny DM, Ongeri F, Perales L, Przytycka TM, Pu LL, Robinson G, Thornton RL, Saada N, Scherer SE, Smith HE, Vinson C, Warner CB, Worley KC, Wu YQ, Zou X, Cherbas P, Kellis M, Eisen MB, Piano F, Kionte K, Fitch DH, Sternberg PW, Cutter AD, Duff MO, Hoskins RA, Graveley BR, Gibbs RA, Bickel PJ, Kopp A, Carninci P, Celniker SE, Oliver B, Richards S. Comparative validation of the D. melanogaster modENCODE transcriptome annotation. Genome Res. 2014 Jul 1;24(7):1209–1223. doi: 10.1101/gr.159384.113. [DOI]

Drosophila 12 Genomes Consortium, Clark, A.G., Eisen, M.B., Smith, D.R., Bergman, C.M., Oliver, B., …, MacCallum, I. 2007. Evolution of genes and genomes on the Drosophila phylogeny. Nature 450(7167): 203--218. PMID: 17994087

Duda, O. 1924 Revision der europäischen u. grönländischen sowie einiger sudostasiat. Arten der Gattung Piophila Fallén (Dipteren) [part]. Konowia 3:97-113 [1924.07.10]

Gonnet, G. H., Cohen, M. A., & Benner, S. A. 1992. Exhaustive matching of the entire protein sequence database. Science (New York, N.Y.), 256(5062):1443–1445. 10.1126/science.1604319. PMID: 1604319

Gramates LS, Agapite J, Attrill H, Calvi BR, Crosby MA, dos Santos G, et al., Lovato. 2022. FlyBase: a guided tour of highlighted features. Genetics 220:10.1093/genetics/iyac035. 10.1093/genetics/iyac035

Grewal SS. 2009. Insulin/TOR signaling in growth and homeostasis: A view from the fly world. Int J Biochem Cell Biology, 41(5):1006–1010, 2009, doi: 10.1016/j.biocel.2008.10.010. PMID: 18992839

Grönke S, Clarke DF, Broughton S, Andrews TD, Partridge L. 2010. Molecular evolution and functional characterization of Drosophila insulin-like peptides. PLoS Genet. 6(2):e1000857. PMID: 20195512

Gruys ML, O’Brien J, Koehler AC, Almazan A, Opperman K, Sterne-Marr R, et al., Reed LK. 2025. Gene model for the ortholog of Ilp3 in Drosophila ananassae. microPublication Biology. 10.17912/micropub.biology.000958.

Gruys ML, Sharp MA, Lill Z, Xiong C, Hark AT, Youngblom JJ, Rele CP, Reed LK. 2025. Gene model for the ortholog of Glys in Drosophila simulans. microPublication Biology. 10.17912/micropub.biology.001168

Hietakangas V and S. M. Cohen. 2009. Regulation of Tissue Growth through Nutrient Sensing. Genetics, 43(1):389–410, doi: 10.1146/annurev-genet-102108-134815. PMID: 19694515

Ikeya T, Galic M, Belawat P, Nairz K, Hafen E. 2002. Nutrient-dependent expression of insulin-like peptides from neuroendocrine cells in the CNS contributes to growth regulation in Drosophila. Curr Biol. 12(15):1293–1300. PMID: 12176357.

Jenkins VK, Larkin A, Thurmond J, FlyBase Consortium. 2022. Using FlyBase: A Database of Drosophila Genes and Genetics. Methods Mol Biol .2540: 1–34. PubMed

Kent, W. J., Sugnet, C. W., Furey, T. S., Roskin, K. M., Pringle, T. H., Zahler, A. M., Haussler, D. 2002. The Human Genome Browser at UCSC. Genome Res. 12:996–1006. PMID: 12045153

Larkin A, Marygold SJ, Antonazzo G, Attrill H, dos Santos G, Garapati PV, Goodman JL, Gramates LS, Millburn G, Strelets VB, Tabone CJ, and Thurmond J and the FlyBase Consortium. 2021. FlyBase: updates to the Drosophila melanogaster knowledge base. Nucleic Acids Res. 49(D1):D899–D907. PMID: 33219682

Morgan A., Kiser C.A., Bronson I., Lin H., Guillette N., McMahon R., Kennell J.A., Long L.J., Reed L.K., Rele C.P. 2022. Drosophila eugracilis — Akt, microPublication Biology. 10.17912/micropub.biology.000544

Mudge JM, Harrow J. 2016. The state of play in higher eukaryote gene annotation. Nat Rev Genet 17(12):758-772. PubMed 10.1038/nrg.2016.119

Myers, A., Hoffman, A., Natysin, M., Arsham, A. M., Stamm, J., Thompson, J. S., Rele, C. P., & Reed, L. K. 2024. Gene model for the ortholog Myc in Drosophila ananassae. microPublication biology. 10.17912/micropub.biology.000856. 10.17912/micropub.biology.000856

Nässel DR, Broeck JV. 2016. Insulin/IGF Signaling in Drosophila and Other Insects: Factors That Regulate Production, Release and Post-Release Action of the Insulin-like Peptides. Cell Mol Life Sci. 73(2):271–290. doi.org/10.1007/s00018-015-2063-3 PMID: 26472340.

Navarro Gonzalez, J., Zweig, A. S., Speir, M. L., Schmelter, D., Rosenbloom, K. R., Raney, B. J., et al. 2021. The UCSC Genome Browser database: 2021 update. Nucleic Acids Research. 49(1), 1046–1057. PMID: 33221922

Pélandakis, M. and Solignac, M. 1993. Molecular phylogeny of Drosophila based on ribosomal RNA sequences. J Mol Evol. 37(5):525–43. doi: 10.1007/BF00160433 PMID: 8283482

Raney BJ, Dreszer TR, Barber GP, Clawson H, Fujita PA, Wang T, Nguyen N, Paten B, Zweig AS, Karolchik D, Kent WJ. 2014. Track data hubs enable visualization of user-defined genome-wide annotations on the UCSC Genome Browser. Bioinformatics. 30(7):1003–5. PMID: 24227676

Rele CP, Sandlin KM, Leung W, Reed LK. 2023. Manual Annotation of Genes within Drosophila Species: the Genomics Education Partnership protocol. F1000Res. 13;11:1579. doi: 10.12688/f1000research.126839.3. PMID: 37854289; PMCID: PMC10579860.

Tello-Ruiz MK, Marco CF, Hsu FM, Khangura RS, Qiao P, Sapkota S, et al., Micklos DA. 2019. Double triage to identify poorly annotated genes in maize: The missing link in community curation. PLoS One 14(10):e0224086. PubMed 10.1371/journal.pone.0224086

Wigby S, Slack C, Grönke S, Martinez P, Calboli FC, Chapman T, Partridge L. 2011. Insulin signaling regulates remating in female Drosophila. Proc Biol Sci. 278(1704):424-431. doi.org/10.1098/rspb.2010.1390 PMID: 20739318

Yamaguchi ST, Tomita J, Kume K. 2022. Insulin signaling in clock neurons regulates sleep in Drosophila. Biochem. Biophys. Res. Commun. 591:44-49. doi.org/10.1016/j.bbrc.2021.12.100 PMID: 34998032

